# Land-use influence on benthic algal assemblage structure and biomass in Afrotropical headwater streams

**DOI:** 10.64898/2026.07.24.740510

**Authors:** Sheila N. Rioba, Jacob O. Iteba, Gideon L. Kuluo, Suzanne R. Jacobs, Lutz Breuer, Frank O. Masese

## Abstract

Benthic algae integrate catchment-and reach-scale pressures in streams. We assessed land-use effects on benthic algal assemblage structure, chlorophyll-a, ash-free dry mass (AFDM), and autotrophic index across 24 headwater streams in the Sondu–Miriu River basin, Kenya, and explored wet–dry patterns in two monitoring streams. We sampled across four land-use categories: natural forest (NF), tea and tree plantations (TTP), smallholder agriculture (SHA), and smallholder tea (SHT). Synoptic sampling occurred in August 2025, and one NF stream and one SHA stream were sampled five times between November 2024 and May 2025. Shannon and Simpson diversity were highest in NF streams and lowest in SHT streams. Chlorophyll-a and AFDM were highest in SHA streams; chlorophyll-a was lowest in SHT, while AFDM and autotrophic index were lowest in NF. Relative abundance patterns showed that SHA streams were dominated by *Lyngbya* and *Stigeoclonium*, whereas SHT streams were dominated mainly by *Stigeoclonium*; NF streams supported persistent diatom-rich assemblages. Non-metric multidimensional scaling showed land-use separation of benthic algal assemblages, while redundancy analysis indicated weak, non-significant evidence of environmental associations in the synoptic dataset. Composition and biomass metrics tracked land-use gradients and wet–dry patterns, supporting periphyton-based bioassessment for monitoring disturbance in Afrotropical headwater streams.

## Introduction

Land-use expansion and intensification in headwater catchments can encroach on riparian corridors, reduce riparian cover integrity and increase hydrological connectivity between hillslopes and stream channels, altering the physical and biogeochemical conditions of headwater streams (Allan, 2004; Gomi et al., 2002; Jackson et al., 2022). In mixed land-use mosaics, which are typical of many tropical catchments, the conversion of forested riparian zones to agriculture, tea cultivation and settlement can act as a cumulative stressor by opening the canopy cover, increasing incident radiation and stream temperature, reducing dissolved oxygen, and increasing sediment and dissolved-solute inputs (Poole & Berman, 2001; Sweeney et al., 2004; Stevenson & Rollins, 2017; Fugère & Masese, 2025; Masese et al., 2024). Because headwater streams are intrinsically linked to their surrounding catchments, such changes can rapidly alter in-stream habitat conditions and restructure benthic primary-producer assemblages, with consequences for basal energy supply and food web functioning in stream ecosystems (Douglas et al., 2005; Allan & Castillo, 2007; Guo et al., 2016).

Conventional assessment of streams and rivers traditionally relies on physical and chemical variables, which only reflect conditions at the time of sampling (Farrell-Poe, 2005; Omer, 2019; Masese et al., 2023). Common indicators include temperature, pH, dissolved oxygen, electrical conductivity, total dissolved solids, total suspended solids, turbidity, particulate organic matter and nutrient fractions, which are often useful for distinguishing land-use effects because agricultural and settlement activities can increase nitrogen and phosphorus inputs through fertiliser, manure, soil disturbance and wastewater pathways (Dessie et al., 2024; Loucif & Chenchouni, 2024; Masese et al., 2024). In tropical catchments, these variables can change rapidly between wet and dry periods because rainfall and runoff alter flow pathways, suspended solids, nutrients, dissolved solutes, and dissolved oxygen conditions over short time periods (Alkhadher et al., 2025; Saturday et al., 2025), emphasizing the sensitivity of physical and chemical water quality parameters to the timing of the sampling. Because of this temporal variability, biological monitoring complements physical and chemical assessment by integrating environmental conditions over longer periods and providing ecologically meaningful signals of disturbance (Feio et al., 2021).

Biological communities such as macroinvertebrates are widely used in stream assessment because of their high taxonomic diversity, varying tolerance levels, and relatively long-life cycles that enable the integration of cumulative and synergistic effects of multiple stressors over several months (Masese et al., 2024; Rosenberg & Resh, 1993). However, macroinvertebrate-based bioassessment can be confounded by natural spatial heterogeneity and seasonal shifts in community composition and abundance, making it difficult to assess ecosystem status using a single index in isolation (Masese et al., 2023; Iteba et al., 2026). Because algal assemblage composition develops over shorter windows than macroinvertebrate assemblages, algae-based indices can capture more recent environmental change while still reflecting conditions beyond the point at which physical and chemical measurements are taken (Stevenson & Rollins, 2017). As primary producers, algae or periphyton are among the first biological groups to respond directly to shifts in light availability and nutrient concentrations, serving as a vital functional complement to the structural indicators provided by macroinvertebrates and other bioindicators (Dalu et al., 2025; Mpopetsi et al., 2025; Masese et al., 2025).

Benthic algae or periphyton are useful bioindicators because they are ubiquitous primary producers, have short generation times, and respond to changes in the physical and chemical conditions of the aquatic environment (Jin & Pan, 2024; Méndez-Zambrano et al., 2024). In streams and rivers, their sensitivity to disturbance can be amplified by the “open canopy” effect, which increases light availability, thereby stimulating autochthonous primary production (Benstead & Pringle, 2004; Fugère et al., 2018; Nansumbi et al., 2025). These changes favour shifts in assemblage structure from shade-tolerant diatom-rich assemblages towards more opportunistic cyanobacterial and filamentous green algal taxa under suitable nutrient and habitat conditions (Bixby et al., 2009; Tromboni et al., 2019; Allan et al., 2021). Apart from shifts in assemblage structure, benthic algae provide other indices of stream disturbance such as chlorophyll-*a* concentration, ash-free dry mass (AFDM) and autotrophic index. Chlorophyll-*a* serves as a proxy for autotrophic biomass, whereas AFDM approximates total organic biofilm mass; their ratio (AFDM/Chl-*a*), expressed as the autotrophic index, can help distinguish autotrophic from heterotrophic dominance under contrasting disturbance regimes (Weber, 1973; Myers, 2014). However, compared with other biological communities such as macroinvertebrates and fish, fewer studies have quantified how land-use gradients translate into shifts in benthic algal assemblage structure and biomass in Afrotropical headwater streams (Bixby et al., 2009; Walsh & Wepener, 2009; Bere et al., 2013; Mangadze et al., 2015; Shibabaw et al., 2021; Dalu et al., 2022). This gap necessitates studies that link land use, physical and chemical gradients, algal assemblage composition and biomass responses to strengthen the use of benthic algae in ecological assessment of Afrotropical headwater streams.

The Sondu–Miriu River basin provides a suitable setting for evaluating the effects of land-use change on benthic algal assemblages because its headwaters drain the Mau Forest Complex, East Africa’s largest remaining tropical Afromontane forest and water tower, before flowing through landscapes that differ in the extent and intensity of forest conversion, tea cultivation and smallholder agriculture (Masese et al., 2017; Jacobs et al., 2018; Stenfert Kroese et al., 2020). This land-use gradient is expected to alter riparian canopy cover, stream temperature, sediment delivery, turbidity, dissolved solutes and oxygen conditions (Guzha et al., 2018; Stenfert Kroese et al., 2020; Wanderi et al., 2022), all of which can influence primary production, benthic algal biomass and assemblage composition (Benstead & Pringle, 2004; Fugère et al., 2018). However, few studies in Afrotropical headwater systems have explicitly quantified how such land-use gradients, together with wet–dry seasonal variation, structure benthic algal assemblages and biomass.

In this study, we investigated benthic algal assemblage structure and periphyton biomass proxies (chlorophyll-a, AFDM, and autotrophic index) as indicators of ecological condition across headwater streams draining contrasting land-use types in the Sondu–Miriu River Basin. The synoptic survey across 24 streams was used to assess land-use effects, while monthly monitoring of one representative natural forest (NF) stream and one representative smallholder agriculture (SHA) stream was used to capture wet–dry seasonal variations. The objectives were to (i) determine whether benthic algal assemblage structure and periphyton biomass differ among land-use types, (ii) assess seasonal variation in algal assemblage structure and biomass in the representative NF and SHA monitoring streams, (iii) identify the physical and chemical variables associated with variation in algal assemblage structure and biomass, and (iv) identify algal taxa associated with land-use categories across the 24 headwater streams and with seasonal variation in the two representative NF and SHA monitoring streams. We hypothesized that (H1) algal assemblages and periphyton biomass proxies would differ among land-use types, with shifts toward disturbance-tolerant, opportunistic taxa and higher chlorophyll-*a* and AFDM in more open-canopy, disturbed reaches; (H2) the representative NF and SHA monitoring streams would show wet–dry seasonal variation in algal assemblage structure and biomass; (H3) variation in benthic algal assemblage structure and periphyton biomass would be associated with land-use-related gradients, with disturbed streams showing higher temperature, turbidity, TSS, EC and TDS, and forested streams showing higher dissolved oxygen; and (H4) the relative abundance of dominant algal genera would differ among land-use categories, with some genera showing contrasting associations with either shaded, less-disturbed forested conditions or agriculturally influenced streams, enabling their future evaluation as indicators of land-use change in tropical headwater streams.

## Materials and Methods

### Study area

This study was carried out in headwater streams within the Sondu Miriu River Basin (SMRB), Kenya. The sampled headwaters drain the south-western block of the Mau Forest Complex, a key Afromontane forest system that functions as a major national “water tower” in Kenya (UNEP, 2008). The basin experiences a bimodal rainfall regime, with the long rains typically occurring from April to July and the short rains from October to December (Koech et al., 2022). The driest months are generally January and February. Human pressure is substantial, with population density around 300 persons km□², and ongoing land transformation has reduced native vegetation cover while increasing sediment and nutrient loading in streams through conversion of native forest to smallholder farms and grazing areas (Jacobs et al., 2017; Stenfert Kroese et al., 2020).

### Study design

To assess land-use influences on headwater stream condition and benthic algal assemblages, study streams in the Sondu–Miriu headwaters were grouped into four land-use categories: natural forest (NF), smallholder agriculture (SHA), smallholder tea (SHT), and tea–tree plantation (TTP). These categories reflect the dominant catchment and riparian land cover around each stream and formed the basis for site classification and land-use comparison in this study (Jacobs et al., 2017; Jacobs et al., 2018). Headwater streams draining catchments dominated by each class were identified, and six independent streams were selected per category, giving a total of 24 study streams (Fig. 1). For the spatial land-use comparison, one synoptic sampling campaign was conducted across 24 streams in August 2025, representing a post-long-rains sampling period. To examine seasonal patterns in water quality and benthic algal communities, two long-term monitoring stream sites, one in NF land use (NF-1) and another in SHA land use (SHA-14), were sampled during five monitoring events between November 2024 and May 2025. Wet-season sampling months were November 2024, April 2025 and May 2025, while dry-season sampling months were January and February 2025.

**Fig. 1.**
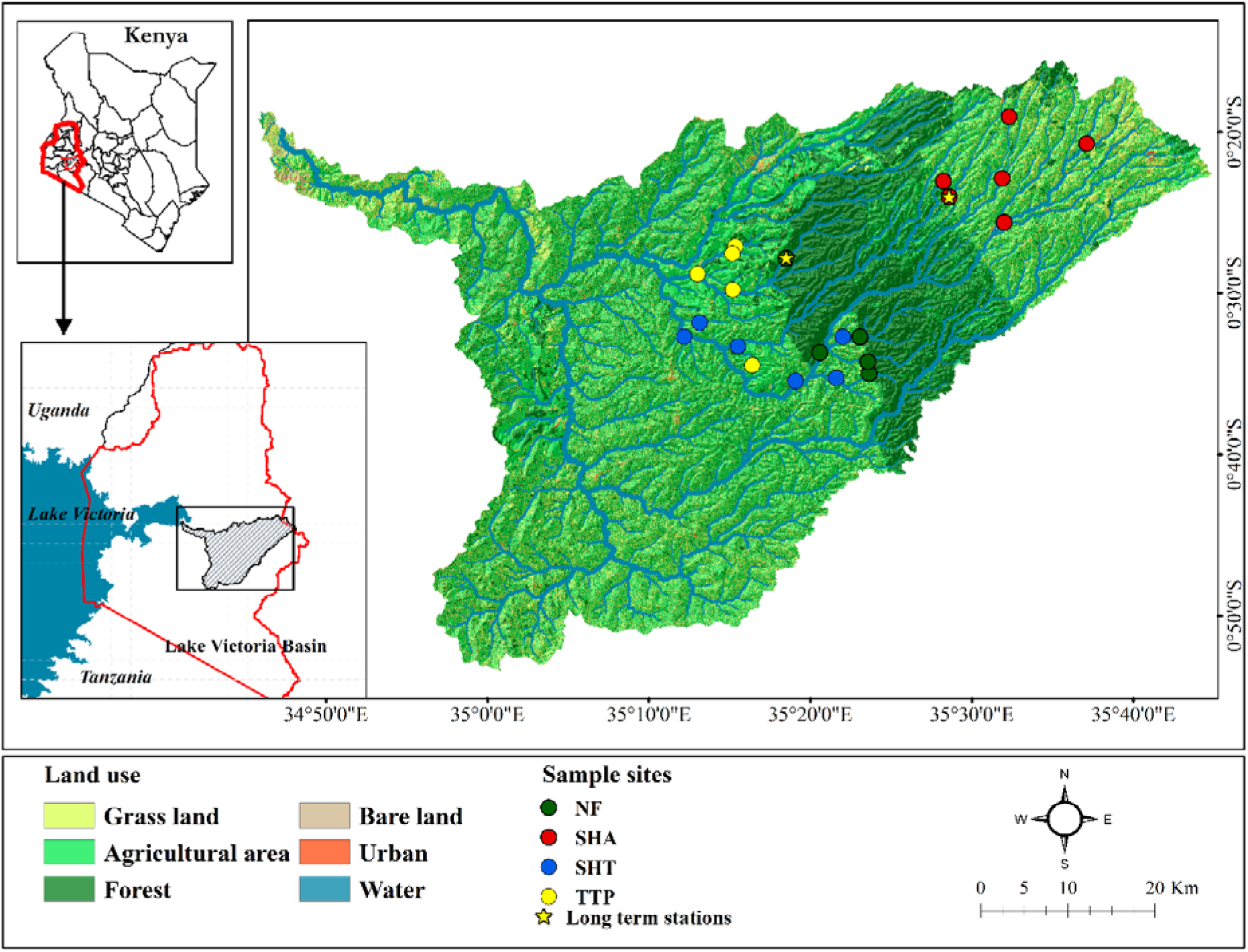
Maps showing the study area location along with sites sampled. Sampling sites are grouped by land-use category: Natural Forest (NF), Smallholder Agriculture (SHA), Smallholder Tea (SHT), and Tea and Tree Plantation (TTP). Land-use classes were derived from the 2013 Landsat-based land-use map of Swart (2016), while sub-catchment delineation was based on a 30-m Shuttle Radar Topography Mission digital elevation model (USGS, 2000). Canopy cover data were obtained from Reiner et al. (2023)

Sub-catchments contributing to each sampling site were delineated using a 30 m SRTM digital elevation model (USGS, 2000). The proportional coverage of different land-use types within each sub-catchment was then quantified using a land-use map derived from 2013 Landsat imagery (Swart, 2016). For most streams, the land-use class used to categorize the stream accounted for >50% of the corresponding sub-catchment area. The four land-use categories were defined according to the dominant catchment and riparian land cover around each stream. NF streams represented the least-disturbed condition, with high native forest cover and relatively intact riparian vegetation. SHA streams drained smallholder farming catchments, typically composed of farms below 2 ha, where food crops and small-scale cash crops occur together with scattered woodlots (Jacobs et al., 2017). These catchments are commonly associated with cultivation close to stream channels, fertilizer and manure application, small-scale livestock keeping, domestic water abstraction, livestock watering and footpath access to streams, which can increase bank disturbance, sediment delivery and nutrient inputs. SHT streams drained smallholder areas where tea was the main cash crop, unlike SHA catchments, which had more mixed cropping systems. Because the Swart (2016) land-use map does not separate smallholder agriculture from smallholder tea, smallholder catchments below the forest belt and within the tea-growing elevation zone, below approximately 2000 m a.s.l., were classified as SHT. Both SHA and SHT catchments also included small livestock keeping and domestic use of stream water. TTP streams drained managed plantation catchments with the highest combined cover of tea and trees. These plantations are intensively managed through fertilizer application and rotation cycles, but most streams retained narrow riparian strips with native forest vegetation and adjacent exotic tree stands, often dominated by *Eucalyptus spp.* (Jacobs et al., 2017; Kuluo et al., 2026; Iteba et al., 2026)

### Field sampling

*In situ* water-quality measurements of temperature, dissolved oxygen (DO) concentration and percentage saturation, pH, electrical conductivity (EC), and total dissolved solids (TDS) were recorded using a multiparameter meter (Multi 3420, WTW GmbH, Weilheim, Germany). Turbidity was measured in the field using a portable turbidity meter (HACH LANGE GmbH, Berlin, Germany). To account for within-reach variability, readings were taken in triplicate at representative points along the reach, and measurements were completed prior to periphyton sampling to avoid potential effects of disturbance on water-quality readings. Water samples for nutrient analyses were collected from the main flowing section of the streams using acid-washed high-density polyethylene (HDPE). Samples for dissolved nutrients (soluble reactive phosphorus, SRP; ammonium, NH□□-N; nitrate, NO□□-N; and nitrite, NO□□-N) were filtered on-site using GF/F filters (0.70 μm pore size; 47 mm diameter) and stored in labelled HDPE bottles. In addition, known volumes of stream water were filtered through pre-weighed GF/F filters for determination of total suspended solids (TSS) and particulate organic matter (POM). The filters with retained matter were packaged in aluminum envelopes. Separate unfiltered water samples were collected for total phosphorus (TP) determination. All samples were kept at 4 °C in a cooler box and transported to the laboratory for analysis. In the laboratory, water samples were refrigerated at 4 °C and analysed as soon as possible, while filters for TSS and POM were stored cold until processing.

Benthic algae (periphyton) were collected from a randomly selected ∼100 m stream reach at each sampling site. Within each reach, stones were used as the sampling surface because they provide relatively stable habitat for attached biofilms. Three stones were sampled to form one replicate, and a total of nine stones were sampled to obtain three replicates per stream and represent within-reach variability. To minimize disturbance effects (e.g., re-suspension of fine sediments and downstream transport of dislodged material), sampling proceeded in a standardized manner by working from downstream to upstream, and stones were selected from comparable microhabitats, mainly riffles and runs with similar depth and flow velocity. For each stone, only the upper exposed surface was sampled. The sampling area was marked using adhesive tape and measured with a 30 cm ruler. Because stone size and shape varied, no fixed scraped area was imposed. Where the stone surface was irregular, the marked surface was measured as closely as possible using the same approach. The marked surface was then scrubbed with a clean toothbrush while being rinsed with distilled water so that loosened biofilm material was washed into a clean collection tray. For each replicate, the slurry from three stones was combined, homogenized and transferred into labelled containers. An aliquot of each homogenized replicate suspension was preserved immediately in Lugol’s iodine solution for subsequent laboratory identification and enumeration, while the remaining suspension was retained for chlorophyll-a and AFDM analyses. The three replicate periphyton slurries from each stream were processed separately in the laboratory. For subsequent analyses, replicate-level measurements were averaged to obtain one stream-level value for each variable, and stream was used as the unit of replication for land-use comparisons.

Periphyton biomass was quantified using chlorophyll-*a* and ash-free dry mass (AFDM). Known volumes of each replicate periphyton slurry were filtered through Whatman GF/F filters. Triplicate filters were prepared for chlorophyll-*a* determination and protected from light by wrapping them in aluminum foil. A second batch of triplicate pre-weighed GF/F filters was prepared for AFDM determination. Filters and vials were labelled with site code, land-use class, replicate number and sampling date, stored in a cooler box, and transported to the laboratory for processing. The Lugol-preserved aliquots were later used for microscopic identification, enumeration, and calculation of benthic algal density and community metrics.

### Laboratory analysis

Filtered water samples were analyzed spectrophotometrically following APHA (2005). SRP was determined using the ascorbic acid method, TP was measured after acid persulphate digestion and analyzed using the same method. NH□□–N was determined using the salicylate–isocyanurate method. NO□□–N and NO□□–N were both measured colorimetrically according to APHA (2005), with NO□□–N determined after cadmium reduction. TSS and POM were determined from stream-water samples filtered through pre-weighed Whatman GF/F filters. Filters with embedded sediments were oven-dried at 60 °C for 48 h (or to constant mass) and reweighed to estimate TSS gravimetrically (APHA, 2005). Filters were then combusted at 450 °C for 4 h, cooled in a desiccator, and reweighed. Mass loss on ignition method was used to quantify POM gravimetrically (APHA, 2005).

Periphyton chlorophyll-*a* was extracted from GF/F filters using 90% acetone and quantified spectrophotometrically following standard methods (APHA, 2005). Absorbance was read at 663 nm and corrected for turbidity/colour by subtracting the 750 nm reading (Wetzel & Likens, 1991). Chlorophyll-*a* concentration was calculated according to Lorenzen (1967). Periphyton AFDM was quantified by loss-on-ignition method as an estimate of total biofilm organic matter (Wetzel & Likens, 2000; APHA, 2005; Steinman et al., 2017). Filters retaining periphyton material were dried at 60 °C for 48 h, cooled in a desiccator, and weighed to obtain dry mass. The same filters were then combusted at 450 °C for 4 h, cooled in a desiccator, and reweighed to obtain ash mass. AFDM was calculated as the difference between dry and ash mass and expressed per unit scraped area.

Benthic algal samples preserved in Lugol’s solution were analyzed to determine assemblage composition and density. Before analysis, each sample was mixed thoroughly, and a 0.5 mL aliquot was transferred to a counting chamber and examined under an inverted light microscope at 200× and 400× magnification. Samples were first scanned at 200× to locate and assess larger algal forms, while genus-level identification and enumeration were conducted mainly at 400×. Counting was done systematically across representative fields of view within the chamber, and all algae observed in the selected fields were recorded. Single-celled taxa were counted as individual cells, while colonial and filamentous taxa were converted to cell counts by enumerating the number of cells within colonies and filament fragments. Taxa were identified to the lowest practical taxonomic level possible, mainly genus, using standard algal identification keys and regional references (Needham & Needham, 1962; Botes, 2003; Janse van Vuuren et al., 2006).

Algal density was calculated by scaling the number of counted algal cells recorded in the 0.5 mL aliquot to the total volume of the homogenized replicate suspension and standardizing by the total scraped area of the three stones forming each replicate. Density was calculated as: D = (N × Vt / Va) / A, where N is the number of counted algal cells recorded in the aliquot, Vt is the total volume of the homogenized replicate suspension, Va is the aliquot volume and A is the total scraped area. Density was expressed as cells m□².

### Data analysis

Water physical and chemical variables were screened for normality and homogeneity of variance. They were natural log-transformed [ln(x+ 1)] where necessary to improve normality and reduce heteroscedasticity (Zar, 2010). For multivariate analyses, environmental variables were standardized using z-score transformation to minimize the influence of differing measurement scales and units (Legendre & Legendre, 2012).

Differences in synoptic physical and chemical variables, biomass metrics, and diversity indices among land-use categories were evaluated using one-way analysis of variance (ANOVA) (Underwood, 1997). Pairwise group differences were evaluated using Tukey’s Honestly Significant Difference (HSD) post hoc tests where appropriate (Tukey, 1949). Results were presented as mean ± standard error (SE), and statistical significance was determined at α = 0.05.

Monthly water physical and chemical variables and periphyton biomass data from the NF and SHA monitoring streams were summarized descriptively by season. Dry-season months were January and February 2025, while wet-season months were November 2024, April 2025 and May 2025. For each variable, values were presented as mean ± standard error (SE) for each monitoring stream and season. Because monthly monitoring was conducted in only one NF stream and one SHA stream, these data were not treated as replicated land-use comparisons.

Benthic algal assemblages were analyzed at the genus level, with pooled stream-level data used for community analyses. Diversity metrics, including Shannon diversity, Simpson diversity, and Pielou’s evenness, were calculated using the ‘vegan’ package (Oksanen et al., 2024). Relative abundance plots were generated separately for each land-use category to visualize dominant and rare genera.

For the synoptic land-use comparison, stream was used as the unit of replication. Measurements from stones, subsamples and within-stream replicates were averaged to obtain one value per stream before analysis, giving six independent streams per land-use category and 24 streams in total. For community analyses, genus-level abundance data were aggregated to stream level before calculating relative abundance and before running NMDS, PERMANOVA, PERMDISP, SIMPER and RDA.

Differences in algal community composition were tested using Permutational Multivariate Analysis of Variance (PERMANOVA) based on Bray-Curtis dissimilarities with 9,999 permutations (Bray & Curtis, 1957; Anderson, 2001). The proportion of variation explained by each factor was quantified using the coefficient of determination (R²). Homogeneity of multivariate dispersions among groups was assessed using Permutational Analysis of Multivariate Dispersions (PERMDISP) to determine whether significant PERMANOVA results reflected differences in community composition rather than differences in within-group variability (Anderson, 2006). Similarity Percentage (SIMPER) analysis was subsequently performed to identify the genera contributing most to compositional differences among groups. The contribution of each genus to Bray-Curtis dissimilarity was calculated and ranked according to its percentage contribution to overall community dissimilarity (Clarke, 1993). All community analyses (NMDS, PERMANOVA, PERMDISP, SIMPER and RDA) were performed using the pooled stream-level community matrices.

Principal Component Analysis (PCA) was used to explore spatial variation in physical and chemical conditions among streams and land-use categories (Legendre & Legendre, 2012). Prior to constrained ordination analyses, Detrended Correspondence Analysis (DCA) was performed to estimate gradient lengths and determine the suitability of linear versus unimodal response models (Hill & Gauch, 1980). Because gradient lengths were generally less than three standard deviation units, linear ordination approaches were considered appropriate, and redundancy analysis (RDA) was applied to evaluate relationships between benthic algal assemblages and environmental variables (ter Braak, 1986; Lepš & Šmilauer, 2003). Prior to RDA, environmental variables were screened for multicollinearity using variance inflation factors (VIF), and variables with VIF > 3 were sequentially removed before fitting the final model. All statistical analyses and figures were produced in R version 4.6 (R Core Team, 2026), using the vegan package (Oksanen et al., 2024).

## Results

### Spatial variation in water quality

Significant spatial variation was observed in several physical and chemical variables across the different land-use categories (Table 1; p < 0.05). The SHT streams recorded the highest mean water temperatures (17.52 ± 0.93 °C), whereas NF streams showed the lowest value (14.92 ± 1.08 °C). In contrast, DO exhibited an opposite spatial pattern, with the highest concentrations measured in NF streams (7.56 ± 0.40 mg/L) and the lowest mean values recorded in SHT streams (6.80 ± 0.39 mg/L). The SHA streams consistently recorded the highest mean electrical conductivity (EC; 52.12 ± 17.47 μS/cm) and total dissolved solids (TDS; 53.50 ± 18.22 mg/L) values, while the lowest concentrations for both variables occurred in NF streams (EC: 25.50 ± 1.42 μS/cm; TDS: 25.50 ± 1.38 mg/L). Turbidity, TSS, POM, pH and nutrient variables, including TP, SRP, NO□□–N and NO□□–N, did not differ significantly among land-use categories (Table 1; p > 0.05).

**Table 1.** Means (±SE) variation of physical and chemical variables in streams draining different land-use categories. NF = natural forest, TTP = tea and tree plantation, SHA = smallholder agriculture, and SHT = smallholder tea. Turbidity, temperature, DO = dissolved oxygen, EC = electrical conductivity, TDS = total dissolved solids, TSS = total suspended solids, POM = particulate organic matter, TP = total phosphorus, SRP = soluble reactive phosphorus, NO –N = nitrate-nitrogen, and NO –N = nitrite-nitrogen. F = one-way ANOVA test statistics. Different superscript letters within a row indicate significant differences among land-use categories at p < 0.05.

| Variable | NF | TTP | SHA | SHT | F | p |
| --- | --- | --- | --- | --- | --- | --- |
| TP (mg/L) | 0.17 ± 0.09<sup>a</sup> | 0.03 ± 0.13<sup>a</sup> | 0.1 ± 0.16<sup>a</sup> | 0.18 ± 0.19<sup>a</sup> | 1.328 | 0.293 |
| SRP (μg/L) | 0.13 ± 0.05<sup>a</sup> | 0.12 ± 0.06<sup>a</sup> | 0.16 ± 0.07<sup>a</sup> | 0.02 ± 0.21<sup>a</sup> | 1.650 | 0.210 |
| NO<sub>3</sub><sup>-</sup>-N (mg/L) | 0.4 ± 0.28<sup>a</sup> | 2.85 ± 1.31<sup>a</sup> | 1.77 ± 1.93<sup>a</sup> | 1.04 ± 0.91<sup>a</sup> | 1.767 | 0.186 |
| NO<sub>2</sub><sup>-</sup>-N (mg/L) | 0.14 ± 0.09<sup>a</sup> | 0.25 ± 0.09<sup>a</sup> | 0.16 ± 0.11<sup>a</sup> | 0.35 ± 0.35<sup>a</sup> | 1.390 | 0.275 |
| TSS (mg/L) | 11.97 ± 3.28<sup>a</sup> | 17.64 ± 10.32<sup>a</sup> | 27.03 ± 9.15<sup>a</sup> | 54.13 ± 40.30<sup>a</sup> | 0.843 | 0.486 |
| POM (mg/L) | 4.69 ± 0.96<sup>a</sup> | 4.96 ± 2.5<sup>a</sup> | 8 ± 4.21<sup>a</sup> | 11.12 ± 7.51<sup>a</sup> | 0.603 | 0.621 |
| Turb (NTU) | 9.25 ± 2.48<sup>a</sup> | 17.27 ± 11.12<sup>a</sup> | 40.65 ± 26.74<sup>a</sup> | 42.55 ± 41.44<sup>a</sup> | 2.615 | 0.079 |
| EC (μS/cm) | 25.5 ± 1.42<sup>b</sup> | 46.67 ± 10.33<sup>ab</sup> | 52.12 ± 17.47<sup>a</sup> | 42.95 ± 17.83<sup>ab</sup> | 4.341 | **0.016** |
| TDS (mg/L) | 25.5 ± 1.38<sup>b</sup> | 46.83 ± 10.42<sup>ab</sup> | 53.5 ± 18.22<sup>a</sup> | 43.17 ± 17.7<sup>ab</sup> | 4.542 | **0.014** |
| DO (mg/L) | 7.56 ± 0.4<sup>a</sup> | 7.07 ± 0.35<sup>ab</sup> | 6.98 ± 0.31<sup>ab</sup> | 6.8 ± 0.39<sup>b</sup> | 4.918 | **0.010** |
| pH | 7.14 ± 0.29<sup>a</sup> | 7.20 ± 0.28<sup>a</sup> | 7.33 ± 0.48<sup>a</sup> | 7.17 ± 0.27<sup>a</sup> | 0.073 | 0.974 |
| Temp ($^{\circ}$C) | 14.92 ± 1.08<sup>b</sup> | 16.12 ± 0.64<sup>b</sup> | 15.72 ± 0.64<sup>b</sup> | 17.52 ± 0.93<sup>a</sup> | 9.977 | **<0.001** |

Principal Component Analysis (PCA) revealed spatial separation among sampling reaches based on their physical and chemical characteristics across the different land use categories (Fig. 2). The first two principal components explained 63.1% of the total variation, with PC1 accounting for 39.1% and PC2 explaining 24.0%. PC1 mainly contrasted DO and SRP with temperature, turbidity, TSS, POM, TP and NO□□–N, whereas PC2 was mainly associated with EC, TDS, NO□□–N and pH. The NF streams clustered closely with the DO vector. In contrast, TTP streams were positioned nearer to the SRP and NO –N vectors, reflecting stronger nutrient-related gradients. The SHA streams were mainly separated along gradients of EC and TDS, while SHT streams showed wide dispersion in ordination space and aligned more strongly with TSS, POM, NO□□–N, turbidity, and temperature gradients.

**Fig. 2.**
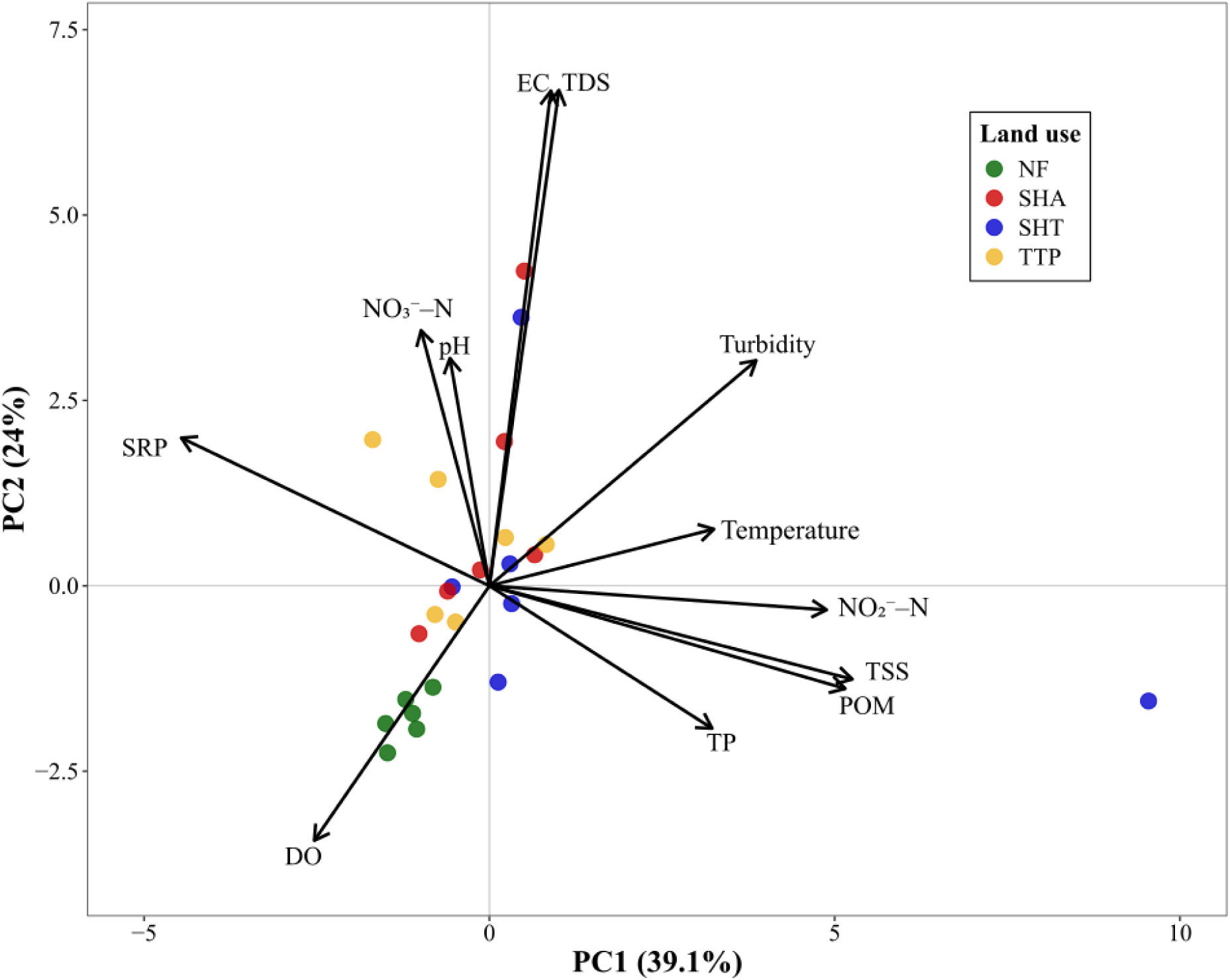
Principal component analysis (PCA) showing variation in water physical and chemical variables among streams draining different land-use categories. Points represent stream samples and vectors represent measured physical and chemical variables. Abbreviations are as defined in Table 1. Axis labels show the percentage of variation explained by each principal component

### Seasonal variation in water quality

Monthly monitoring showed consistent differences in water physical and chemical conditions between the two NF and SHA monitoring streams across dry-and wet-season months (Table 2). DO and DO saturation were higher in the NF stream in both seasons, with the highest values recorded during the dry season. In contrast, the SHA stream had consistently higher EC and TDS than the NF stream. Wet-season months had higher turbidity, TSS and POM than dry-season months in both streams, with the highest values recorded in the SHA stream. Nutrient concentrations also increased during wet-season months, particularly nitrate, nitrite, TP, SRP and NH□□–N. The strongest wet-season increase was observed in the SHA stream, where nitrate, TSS, turbidity and POM were higher than during dry-season months. The monthly data indicate that the SHA stream was characterized by higher dissolved solids, suspended material, particulate organic matter and nutrient concentrations, whereas the NF stream maintained higher oxygen conditions. These patterns are presented as descriptive trends from the two monitored streams.

**Table 2.** Means (±SE) variation of physical and chemical variables in natural forest (NF) and smallholder agriculture (SHA) monitoring streams during dry-season months and wet-season months. Dry-season months = January and February 2025; wet-season months = November 2024, April 2025 and May 2025.

| Parameter | Season | NF | SHA |
| --- | --- | --- | --- |
| DO (mg/L) | Wet | 7.2 ± 0.2 | 6.6 ± 0.2 |
| | Dry | 7.8 ± 0.1 | 7.1 ± 0.0 |
| DO Saturation (%) | Wet | 94.6 ± 1.7 | 85.1 ± 4.5 |
| | Dry | 104.4 ± 4 | 96.6 ± 0.2 |
| EC (μS/cm) | Wet | 26.5 ± 0.6 | 52.8 ± 0.4 |
| | Dry | 28.8 ± 0.4 | 58.6 ± 5.6 |
| NH<sub>4</sub><sup>+</sup>-N (μg/L) | Wet | 0.40 ± 0.09 | 0.48 ± 0.19 |
| | Dry | 0.10 ± 0.02 | 0.10 ± 0.07 |
| NO<sub>3</sub><sup>-</sup>-N (mg/L) | Wet | 0.59 ± 0.23 | 0.73 ± 0.24 |
| | Dry | 0.05 ± 0.03 | 0.06 ± 0.05 |
| NO<sub>2</sub><sup>-</sup>-N (mg/L) | Wet | 0.82 ± 0.34 | 2.72 ± 1.33 |
| | Dry | 0.06 ± 0.02 | 0.08 ± 0.03 |
| POM (mg/L) | Wet | 19.80 ± 1.95 | 37.27 ± 7.69 |
| | Dry | 8.18 ± 2.08 | 18.82 ± 1.98 |
| SRP (μg/L) | Wet | 0.31 ± 0.14 | 0.36 ± 0.15 |
| | Dry | 0.01 ± 0.00 | 0.03 ± 0.02 |
| TDS (mg/L) | Wet | 28.0 ± 1.2 | 52.1 ± 0.5 |
| | Dry | 33.0 ± 1.1 | 60.5 ± 2.5 |
| TP (mg/L) | Wet | 0.17 ± 0.07 | 0.27 ± 0.09 |
| | Dry | 0.01 ± 0.00 | 0.06 ± 0.02 |
| TSS (mg/L) | Wet | 26.0 ± 0.2 | 76.6 ± 13.1 |
| | Dry | 13.8 ± 1.1 | 44.4 ± 2.5 |
| Temperature ($^{\circ}$C) | Wet | 15.1 ± 0.1 | 16.1 ± 0.3 |
| | Dry | 14.4 ± 0.3 | 15.4 ± 0.1 |
| Turbidity (NTU) | Wet | 16.7 ± 2.4 | 59.3 ± 16.5 |
| | Dry | 7.3 ± 1.7 | 17.55 ± 2.8 |

Principal Component Analysis (PCA) showed separation between NF and SHA streams during both wet and dry seasons (Fig. 3). In the wet season, the first two principal components explained 92.2% of the total variation, with PC1 accounting for 67.7% and PC2 explaining 24.5%. NF samples were positioned on the positive side of PC1 and clustered closer to DO. The SHA samples were positioned on the negative side of PC1 and were more closely aligned with EC, TDS, TSS, POM, TP and SRP.

**Fig. 3.**
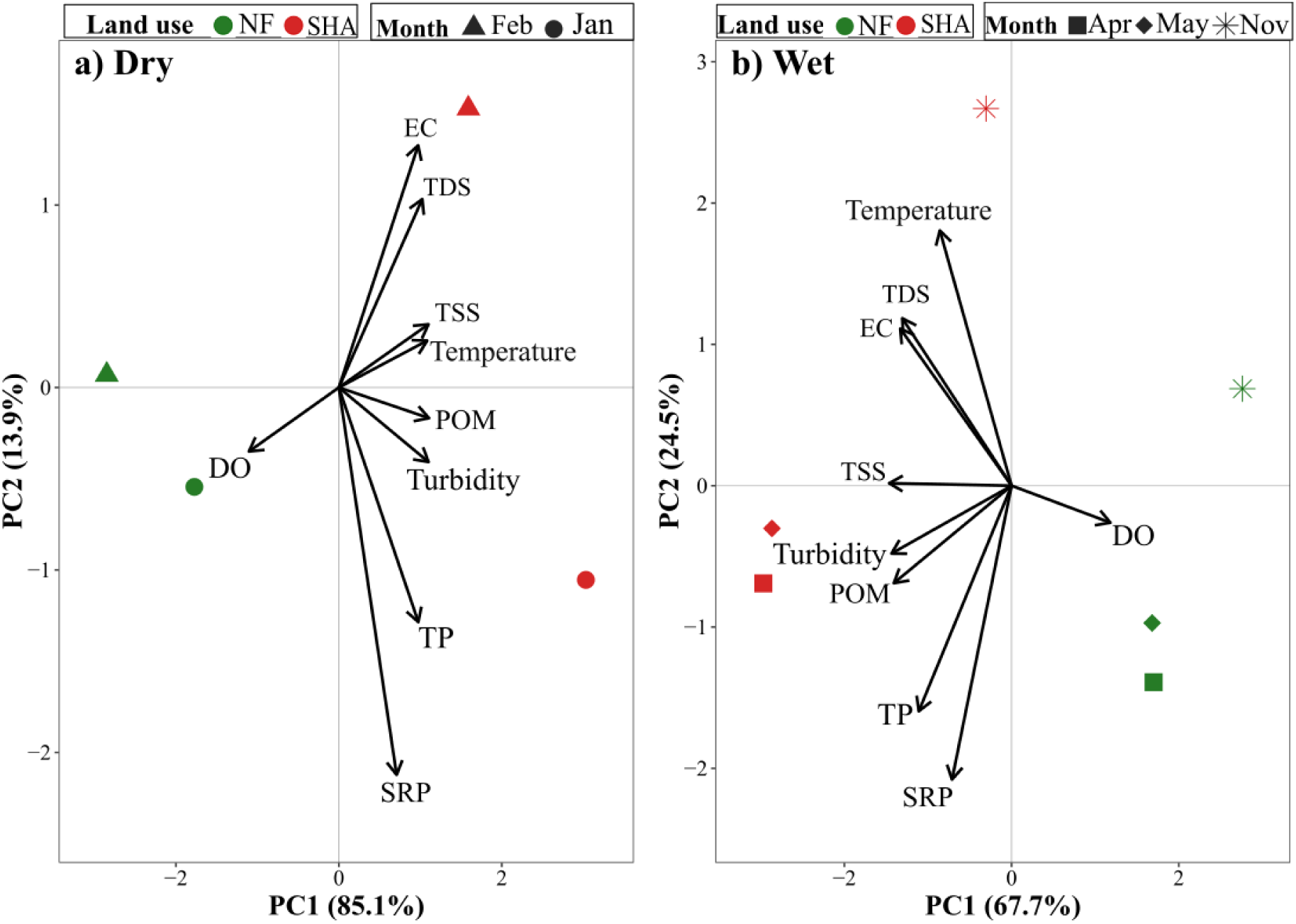
Principal component analysis (PCA) showing variation in physical and chemical variables measured in natural forest (NF) and smallholder agriculture (SHA) monitoring streams during the dry-season months, January–February 2025, and wet-season months, November 2024 and April–May 2025. Abbreviations are as defined in Table 1. Points represent stream samples, and vectors represent the measured physical and chemical variables

In the dry season, the first two principal components explained 99% of the total variation, with PC1 accounting for 85.1% and PC2 explaining 13.9%. PC1 mainly represented gradients associated with EC, TDS, TSS, POM, TP and SRP on the positive side of the axis, whereas DO was positioned toward the negative side of PC1. PC2 was more strongly associated with EC, TDS and TSS in the upper part of the ordination, while TP and SRP were positioned toward the lower part. The SHA samples were generally positioned on the positive side of PC1 and were closer to EC, TDS, TSS, POM, TP and SRP. The NF samples were separated from SHA samples and were positioned closer to the DO vector.

### Spatial variation in benthic algal community composition

Benthic algal assemblage structure showed differences among land-use categories. The NF streams were characterized by diatom-rich assemblages such as *Diatoma*, *Cyclotella*, and *Eunotia.* TTP streams had a high contribution of the filamentous green alga *Stigeoclonium*, together with diatom genera such as *Diatoma*, *Synedra* and *Gomphonema*. The SHA streams were dominated mainly by *Stigeoclonium* and the cyanobacterium *Lyngbya* with additional contributions from *Synedra*, *Microcystis*, *Oscillatoria* and *Navicula*. SHT streams showed the strongest dominance by *Stigeoclonium*, while *Phormidium*, *Synedra*, *Melosira* and *Navicula* occurred at lower relative abundances (Fig. 4). Taxon occurrence across land-use categories is shown in Table 3.

**Fig. 4.**
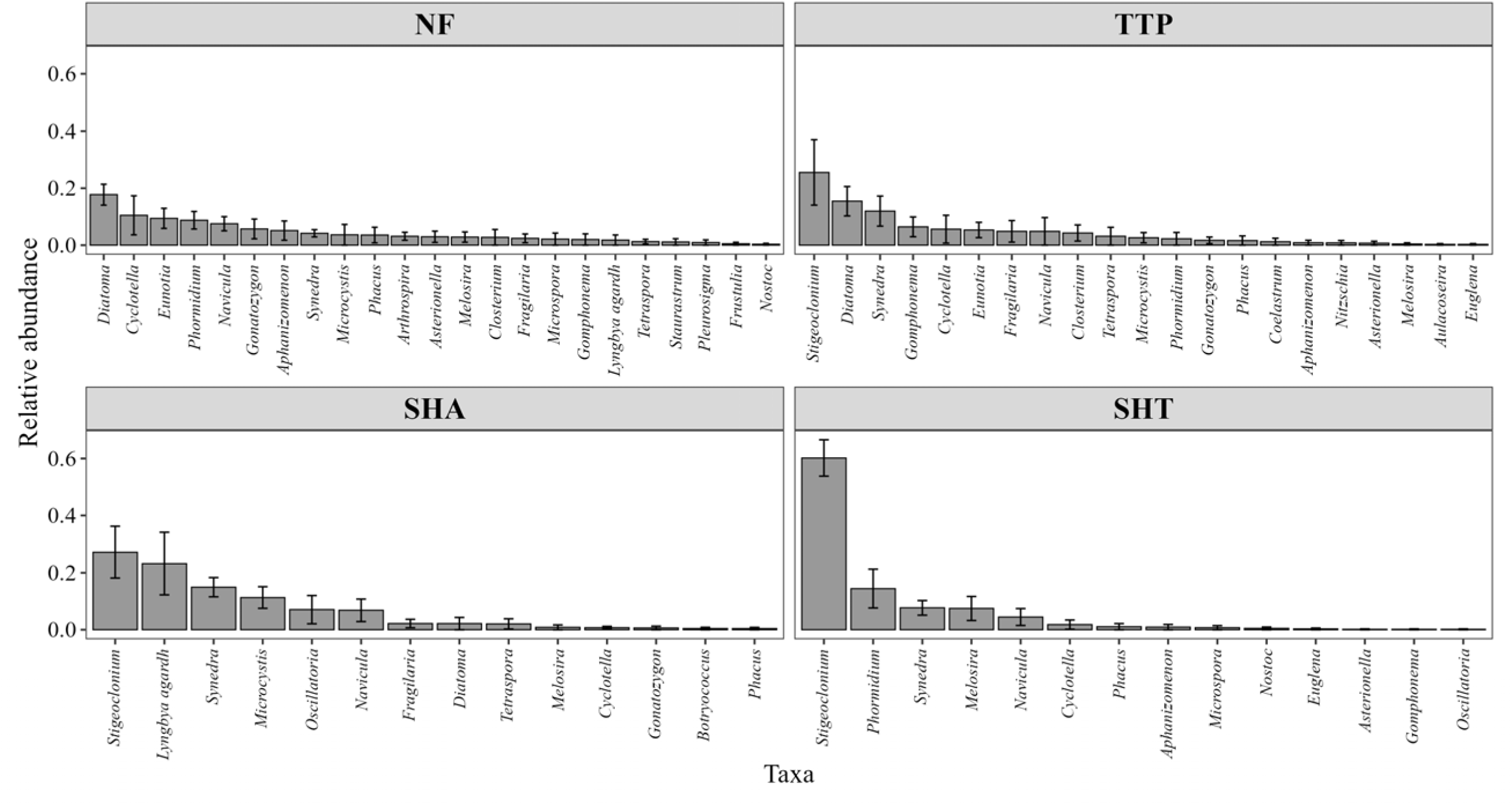
Relative abundance of benthic algal genera within each land-use category. Bars represent the proportional contribution of each genus to total algal density within each land use. NF = Natural Forest, SHA = Smallholder Agriculture, SHT = Smallholder Tea, TTP = Tea and Tree Plantation. Panels correspond to land-use types, and only genera present in each land use are shown. Taxa are ordered from highest to lowest relative abundance from left to right. Error bars represent standard error among streams

**Table 3.** Presence–absence distribution of benthic algal genera across streams draining different land-use categories. “+” indicates presence and “–” indicates absence. NF = natural forest; SHA = smallholder agriculture; SHT = smallholder tea; TTP = tea and tree plantation.

| *Taxa* | NF | SHA | SHT | TTP |
| --- | --- | --- | --- | --- |
| *Ankistrodesmus* | – | – | + | + |
| *Aphanizomenon* | + | – | + | + |
| *Arthrospira* | + | – | – | – |
| *Asterionella* | + | – | + | + |
| *Aulacoseira* | – | – | – | + |
| *Botryococcus* | – | + | – | – |
| *Closterium* | + | – | – | + |
| *Coelastrum* | – | – | – | + |
| *Cyclotella* | + | + | + | + |
| *Diatoma* | + | + | – | + |
| *Euglena* | – | – | + | + |
| *Eunotia* | + | – | – | + |
| *Fragilaria* | + | + | – | + |
| *Frustulia* | + | – | – | – |
| *Gomphonema* | + | – | + | + |
| *Gonatozygon* | + | + | – | + |
| *Lyngbya* | + | + | – | – |
| *Melosira* | + | + | + | + |
| *Merismopedia* | – | – | – | – |
| *Microcystis* | + | + | – | + |
| *Microspora* | + | – | + | – |
| *Navicula* | + | + | + | + |
| *Nitzschia* | – | – | – | + |
| *Nostoc* | + | – | + | – |
| *Oscillatoria* | – | + | + | – |
| *Phacus* | + | + | + | + |
| *Phormidium* | + | – | + | + |
| *Pleurosigma* | + | – | – | – |
| *Staurastrum* | + | – | – | – |
| *Stigeoclonium* | – | + | + | + |
| *Synedra* | + | + | + | + |
| *Tetraspora* | + | + | – | + |

### Seasonal variation in benthic algal community composition

Benthic algal composition differed between the NF and SHA monitoring streams, while wet-and dry-season samples showed descriptive variation in assemblage composition (Fig. 5). The NF stream remained diatom-rich in both seasons, with *Diatoma*, *Eunotia*, *Cyclotella*, *Fragilaria*, *Navicula* and *Nitzschia* contributing most of the assemblage. In contrast, the SHA stream was characterized by higher relative abundances of cyanobacterium *Lyngbya* and filamentous green alga *Stigeoclonium* in both seasons, with additional contributions from *Oscillatoria*, *Gomphonema*, *Nitzschia* and *Microcystis* (Fig. 5). Monthly occurrence patterns across the two monitored streams are summarized in Table 4.

**Fig. 5.**
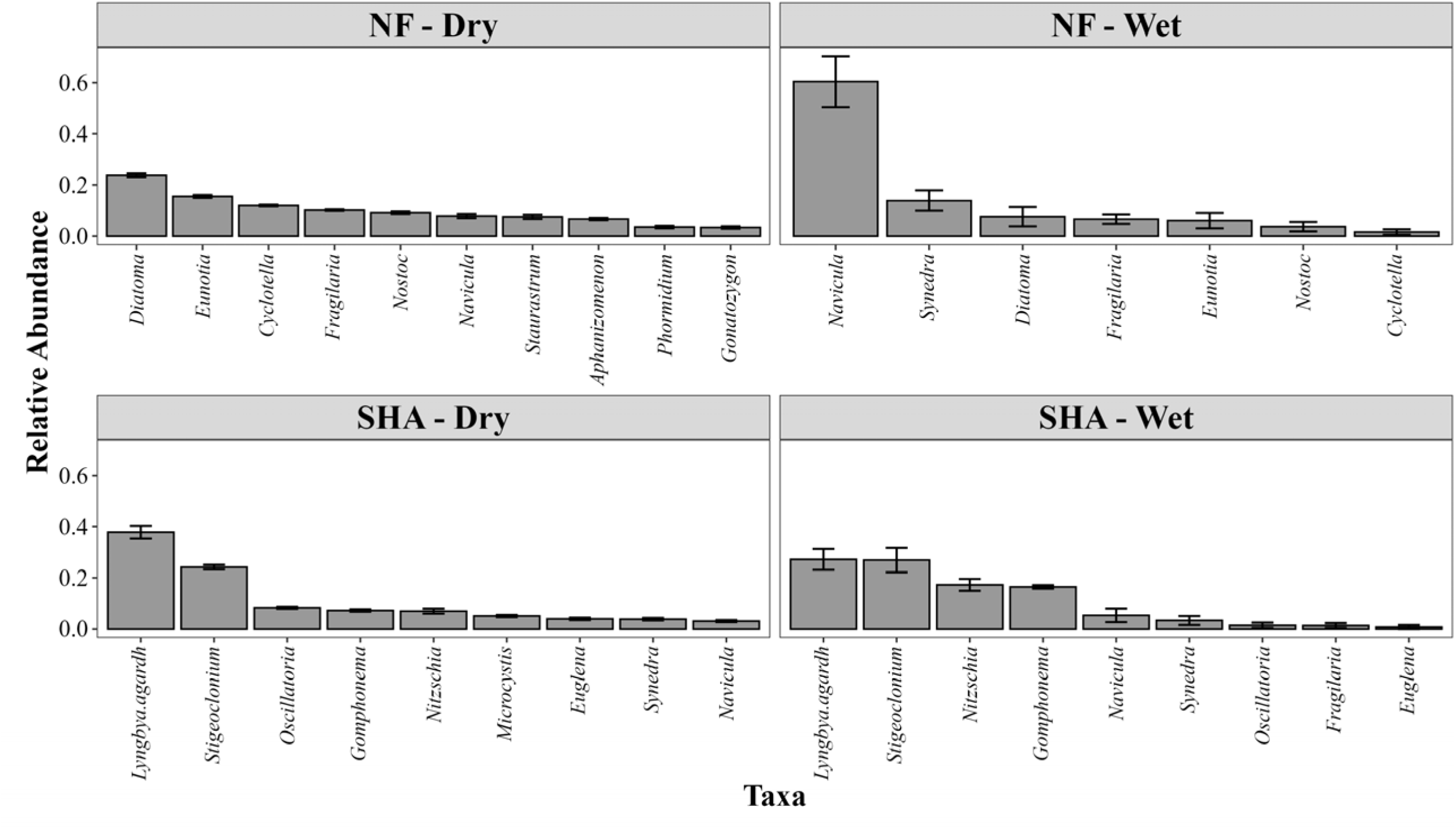
Relative abundance of dominant benthic algal genera in natural forest (NF) and smallholder agriculture (SHA) monitoring streams during dry and wet seasons. Dry = January–February 2025, Wet = November 2024, April–May 2025. Bars represent mean relative abundance, and error bars indicate standard error

**Table 4.** Presence–absence distribution of benthic algal genera recorded during monthly monitoring of natural forest and smallholder agriculture streams from November 2024 to May 2025. “+” indicates presence and “–” indicates absence. Dry season = January– February 2025; wet season = November 2024 and April–May 2025.

| Taxa | NOV | JAN | FEB | APR | MAY |
| --- | --- | --- | --- | --- | --- |
| *Aphanizomenon* | – | + | + | – | – |
| *Cyclotella* | – | + | + | + | – |
| *Diatoma* | + | + | + | – | – |
| *Euglena* | – | + | + | + | – |
| *Eunotia* | + | + | + | – | – |
| *Fragilaria* | + | + | + | + | + |
| *Gomphonema* | + | + | + | + | + |
| *Gonatozygon* | – | + | + | – | – |
| *Lyngbya* | + | + | + | + | + |
| *Microcystis* | – | + | + | – | – |
| *Navicula* | + | + | + | + | + |
| *Nitzschia* | + | + | + | + | + |
| *Nostoc* | + | + | + | – | – |
| *Oscillatoria* | – | + | + | + | – |
| *Phormidium* | – | + | + | – | – |
| *Staurastrum* | – | + | + | – | – |
| *Stigeoclonium* | + | + | + | + | + |
| *Synedra* | + | + | + | + | + |

### Spatial variation in diversity indices and algal biomass

Diversity indices and biomass measures varied among land-use categories (Table 5). Shannon and Simpson diversity differed significantly among land uses, with the main contrast occurring between NF and SHT streams; SHA and TTP streams showed intermediate diversity and did not differ significantly from either group. Chlorophyll-*a* varied significantly across land uses, with the highest value recorded in SHA streams (6.82 ± 0.37 mg m ²) and the lowest in SHT streams (4.54 ± 0.42 mg m ²; Table 5). The AFDM also differed among land use categories (F = 7.660, p = 0.001), with the highest value recorded in SHA streams; NF, TTP and SHT streams did not differ significantly from one another (Table 5). Autotrophic index also differed significantly among land uses, with SHA streams having significantly higher values than NF streams, whereas TTP and SHT streams were intermediate (Table 5). As an AFDM/chlorophyll-a ratio, higher autotrophic index values indicate more organic or heterotrophic biofilm material relative to algal pigment.

**Table 5.** Means (±SE) variation of benthic algal density, diversity indices, and biomass measures in streams draining different land-use categories. NF = natural forest, TTP = tea and tree plantation, SHA = smallholder agriculture, and SHT = smallholder tea. Chl-a = chlorophyll-a; AFDM = ash-free dry mass. F = ANOVA test statistic. Different superscript letters within a row indicate significant differences among land-use categories at p < 0.05.

| Metric | NF | TTP | SHA | SHT | F | *p* |
| --- | --- | --- | --- | --- | --- | --- |
| Total density (cells/m<sup>2</sup>) | 653520.49 ± 99756.43<sup>a</sup> | 810791.28 ± 201718.57<sup>a</sup> | 932550.74 ± 282169.33<sup>a</sup> | 867439.27 ± 153935.95<sup>a</sup> | 0.369 | 0.7759 |
| Shannon (H') | 1.97 ± 0.11<sup>a</sup> | 1.53 ± 0.15<sup>ab</sup> | 1.41 ± 0.12<sup>ab</sup> | 1.05 ± 0.18<sup>b</sup> | 7.064 | 0.0020** |
| Simpson (1-D) | 0.84 ± 0.02<sup>a</sup> | 0.73 ± 0.04<sup>ab</sup> | 0.69 ± 0.04<sup>ab</sup> | 0.55 ± 0.07<sup>b</sup> | 6.603 | 0.0028** |
| Evenness (H'/lnS) | 0.92 ± 0.01<sup>a</sup> | 0.86 ± 0.04<sup>a</sup> | 0.81 ± 0.04<sup>a</sup> | 0.75 ± 0.06<sup>a</sup> | 3.043 | 0.0530 |
| Chl-a (mg/m<sup>2</sup>) | 6.24 ± 0.85<sup>ab</sup> | 4.61 ± 0.51<sup>ab</sup> | 6.82 ± 0.37<sup>a</sup> | 4.54 ± 0.42<sup>b</sup> | 4.139 | 0.0196* |
| AFDM (mg/m<sup>2</sup>) | 666.79 ± 145.64<sup>b</sup> | 838.00 ± 141.53<sup>b</sup> | 1707.27 ± 235.88<sup>a</sup> | 962.56 ± 114.07<sup>b</sup> | 7.660 | 0.0013** |
| Autotrophic Index | 120.65 ± 29.03<sup>b</sup> | 182.66 ± 23.71<sup>ab</sup> | 256.77 ± 39.81<sup>a</sup> | 228.24 ± 38.51<sup>ab</sup> | 3.155 | 0.0474* |

### Seasonal variation in diversity indices and algal biomass

Monthly periphyton biomass metrics varied between the NF and SHA monitoring streams across wet-and dry-season months (Table 6). Chlorophyll-a was higher during dry months than wet months in both streams, increasing from 5.89 ± 1.19 to 30.28 ± 4.23 mg m ² in the NF stream and from 7.91 ± 1.15 to 38.00 ± 10.47 mg m ² in the SHA stream. AFDM also increased during dry months in both streams, from 1117.33 ± 136.43 to 2022.68 ± 198.70 mg m ² in NF and from 1509.66 ± 213.33 to 3971.51 ± 639.62 mg m ² in SHA. The highest AFDM value was recorded in the SHA stream during dry months. During dry-season months, the autotrophic index was higher in the monitored SHA stream (131.34 ± 20.43) than in the monitored NF stream (69.37 ± 4.31).

**Table 6.** Means (±SE) variation of benthic algal biomass measures in natural forest (NF) and smallholder agriculture (SHA) monitoring streams during wet and dry seasons. Chl-a = chlorophyll-a; AFDM = ash-free dry mass.

| Parameter | Season | NF | SHA |
| --- | --- | --- | --- |
| Chlorophyll a (mg/m<sup>2</sup>) | Wet | 5.89 ± 1.19 | 7.91 ± 1.15 |
| Chlorophyll a (mg/m<sup>2</sup>) | Dry | 30.28 ± 4.23 | 38 ± 10.47 |
| AFDM (mg/m<sup>2</sup>) | Wet | 1117.33 ± 136.43 | 1509.66 ± 213.33 |
| AFDM (mg/m<sup>2</sup>) | Dry | 2022.68 ± 198.7 | 3971.51 ± 639.62 |
| Autotrophic Index | Wet | 285.45 ± 62.76 | 193.67 ± 8.63 |
| Autotrophic Index | Dry | 69.37 ± 4.31 | 131.34 ± 20.43 |

### Spatial and seasonal patterns in algal community composition

Non-metric multidimensional scaling (NMDS) ordinations, based on Bray-Curtis dissimilarity matrices, revealed spatial variation in benthic algal assemblage composition across the land use gradient (Fig. 6). The two-dimensional solution reached a goodness-of-fit, providing a reliable representation of the community data (stress = 0.164). NF streams formed a cluster that was separated from other land use categories. In contrast, SHA and SHT streams occupied intermediate positions, while TTP streams exhibited greater dispersion. These observed ordination patterns were statistically corroborated by PERMANOVA, which indicated that land use significantly influenced algal community composition (Pseudo-F = 3.97, R^2^ = 0.373, *p* = 0.0001). Land use accounted for 37.3% of the total variation in benthic algal community structure. Tests for homogeneity of multivariate dispersion were not significant (PERMDISP: F = 1.85, *p* = 0.174), suggesting that the observed differences were attributable to genuine shifts in community composition rather than unequal within-group variability. Pairwise PERMANOVA comparisons, following Benjamini-Hochberg correction, revealed significant differences among most land-use categories. Higher compositional separation occurred between NF and SHT streams (R² = 0.455, adjusted *p* = 0.0048), followed by NF and SHA reaches (R²=0.364, adjusted p = 0.0048). Significant dissimilarities were also detected between SHT and SHA (R²= 0.308), TTP and SHT (R²=0.273), and TTP and SHA streams (R² = 0.196; all adjusted p < 0.05). In contrast, no significant difference in community structure was detected between NF and TTP streams (R²= 0.099, adjusted *p* = 0.356).

**Fig. 6.**
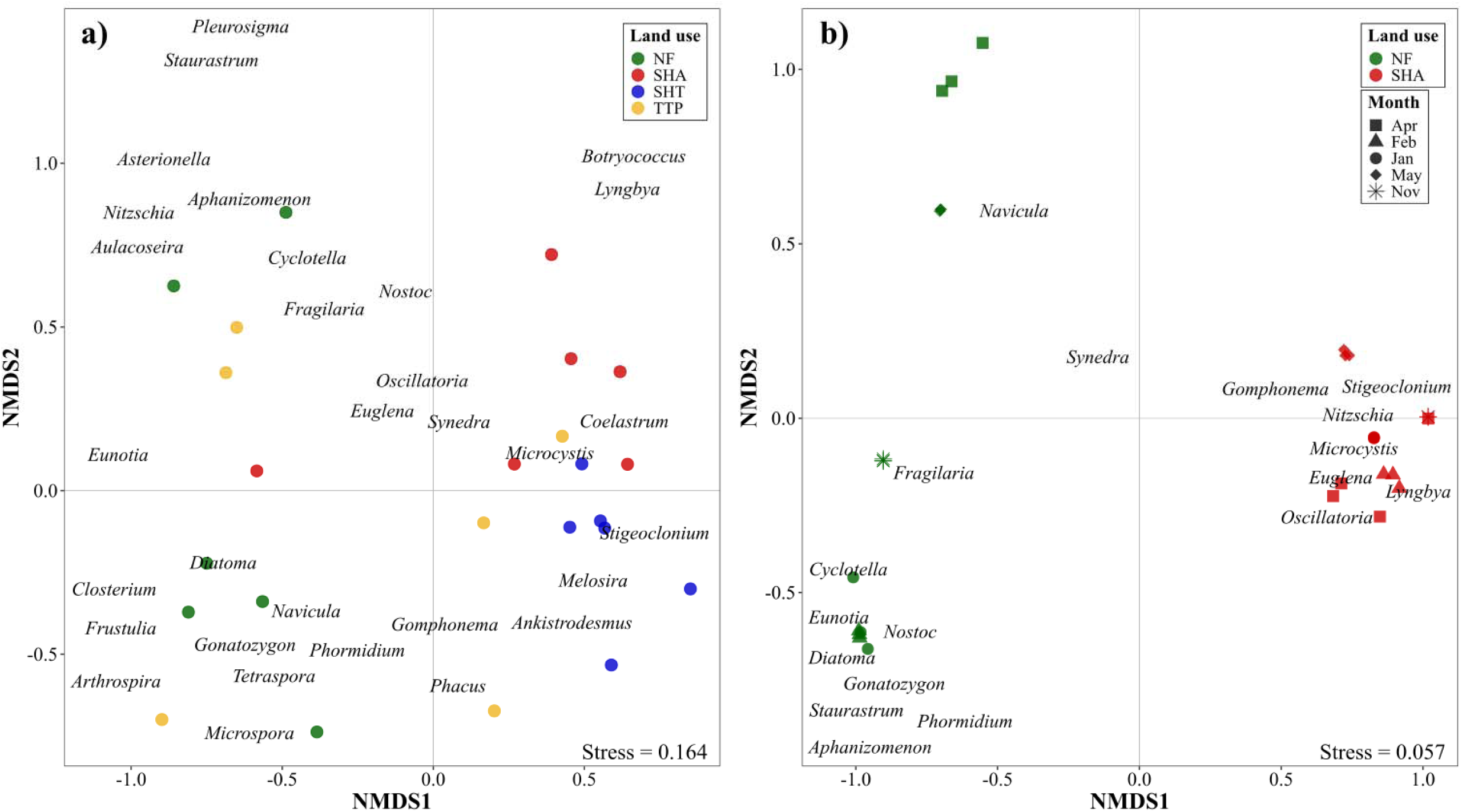
Non-metric multidimensional scaling (NMDS) ordination of benthic algal assemblage composition based on Bray–Curtis dissimilarities for (a) synoptic samples collected across streams draining four land-use categories and (b) monthly monitoring samples collected from natural forest (NF) and smallholder agriculture (SHA) streams. In panel (a), colours represent land-use categories: NF = natural forest, TTP = tea and tree plantation, SHA = smallholder agriculture, and SHT = smallholder tea. In panel (b), colours represent land-use and symbols represent sampling months. Dry season = January–February 2025; wet season = November 2024 and April–May 2025. Taxon names show the position of benthic algal genera in the ordination space. Stress values are shown in each panel

Monthly monitoring NMDS ordination showed that samples from the NF and SHA streams separated mainly by stream/land-use identity, while wet-and dry-season samples overlapped considerably (stress = 0.057; Fig. 6b). PERMANOVA based on Bray-Curtis dissimilarities indicated that differences between the monitored NF and SHA streams explained the largest proportion of the variation in benthic algal community composition (*df = 1, pseudo-F = 30.60, R² = 0.675, p = 0.0011*). Season accounted for a smaller proportion of the variation (*df = 1, pseudo-F = 3.68, R² = 0.081, p = 0.0523*) and was not statistically significant. The interaction between stream and season was significant (*df = 1, pseudo-F = 5.05, R² = 0.111, p = 0.0094*), indicating that seasonal changes in community composition differed between the two monitored streams. Together, stream, season and their interaction explained 86.8% of the total variation in benthic algal community composition, while the remaining 13.2% was attributed to unexplained variation. Because monthly monitoring was conducted in one NF stream and one SHA stream, these community results describe temporal patterns from the two monitored streams.

Similarity percentages (SIMPER) analysis of synoptic data indicated that these compositional differences were attributable mainly to a limited subset of genera (Table 7). Across all pairwise comparisons, *Stigeoclonium*, *Lyngbya*, *Diatoma*, *Cyclotella*, *Phormidium*, *Synedra*, and *Microcystis* contributed most to Bray-Curtis dissimilarity. Higher contribution was observed in the NF-SHT comparison, where *Stigeoclonium* alone accounted for 34.8% of the total dissimilarity, followed by *Phormidium* (10.1%). Differences between NF and SHA streams were associated mainly with higher contributions of *Lyngbya* (16.5%), *Stigeoclonium* (15.5%), and *Microcystis* (5.6%) in SHA streams. Comparisons involving TTP streams were primarily influenced by variation in *Stigeoclonium*, *Diatoma*, *Synedra*, and *Gomphonema*.

**Table 7.** Main benthic algal genera contributing to Bray–Curtis dissimilarity among land-use categories based on SIMPER analysis. Mean densities are presented as log10(cells m ² + 1). Contribution (%), cumulative contribution (%), and permutation p-values are based on the original density matrix.

| Comparison | Genus | Mean density<br>(Group 1) | Mean density<br>(Group 2) | Contribution (%) | Cumulative (%) | p |
| --- | --- | --- | --- | --- | --- | --- |
| NF vs TTP | *Stigeoclonium* | 0.00 | 5.57 | 12.45 | 12.45 | 0.989 |
|  | *Diatoma* | 5.33 | 5.40 | 8.54 | 20.99 | 0.108 |
|  | *Cyclotella* | 5.30 | 4.66 | 7.71 | 28.70 | 0.035 |
|  | *Synedra* | 4.71 | 5.24 | 6.25 | 34.95 | 0.257 |
|  | *Gomphonema* | 4.22 | 5.17 | 5.10 | 40.05 | 0.022 |
| NF vs SHT | *Stigeoclonium* | 0.00 | 5.99 | 34.78 | 34.78 | 0.001 |
|  | *Phormidium* | 4.89 | 5.50 | 10.09 | 44.87 | 0.006 |
|  | *Diatoma* | 5.33 | 0.00 | 7.84 | 52.71 | 0.208 |
|  | *Cyclotella* | 5.30 | 3.99 | 6.10 | 58.81 | 0.222 |
|  | *Eunotia* | 5.11 | 0.00 | 4.40 | 63.21 | 0.032 |
| NF vs SHA | *Lyngbya* | 4.15 | 5.84 | 16.52 | 16.52 | 0.007 |
|  | *Stigeoclonium* | 0.00 | 5.66 | 15.52 | 32.04 | 0.856 |
|  | *Diatoma* | 5.33 | 3.61 | 8.38 | 40.42 | 0.120 |
|  | *Cyclotella* | 5.30 | 4.23 | 6.35 | 46.77 | 0.155 |
|  | *Microcystis* | 4.47 | 5.21 | 5.61 | 52.38 | 0.005 |
| TTP vs SHT | *Stigeoclonium* | 5.57 | 5.99 | 25.35 | 25.35 | 0.017 |
|  | *Phormidium* | 3.65 | 5.50 | 9.47 | 34.82 | 0.019 |
|  | *Diatoma* | 5.40 | 0.00 | 8.08 | 42.90 | 0.200 |
|  | *Synedra* | 5.24 | 5.10 | 5.77 | 48.67 | 0.367 |
|  | *Gomphonema* | 5.17 | 3.36 | 4.29 | 52.96 | 0.117 |
| TTP vs SHA | *Lyngbya* | 0.00 | 5.84 | 16.18 | 16.18 | 0.012 |
|  | *Stigeoclonium* | 5.57 | 5.66 | 15.21 | 31.39 | 0.884 |
|  | *Diatoma* | 5.40 | 3.61 | 8.67 | 40.06 | 0.077 |
|  | *Synedra* | 5.24 | 5.33 | 6.97 | 47.03 | 0.074 |
|  | *Microcystis* | 4.57 | 5.21 | 5.27 | 52.30 | 0.029 |
| SHT vs SHA | *Stigeoclonium* | 5.99 | 5.66 | 19.88 | 19.88 | 0.305 |

| Comparison | Genus | Mean density<br>(Group 1) | Mean density<br>(Group 2) | Contribution (%) | Cumulative (%) | *p* |
| --- | --- | --- | --- | --- | --- | --- |
|  | *Lyngbya* | 0.00 | 5.84 | 14.90 | 34.78 | 0.029 |
|  | *Phormidium* | 5.50 | 0.00 | 8.81 | 43.59 | 0.042 |
|  | *Synedra* | 5.10 | 5.33 | 4.94 | 48.53 | 0.662 |
|  | *Microcystis* | 0.00 | 5.21 | 4.89 | 53.42 | 0.063 |

Similarity percentage (SIMPER) analysis for monthly monitoring showed that differences in community composition were largely attributable to a small number of dominant genera (Table 8). The highest contributions to Bray-Curtis dissimilarity were provided by *Lyngbya* (19.4%) and *Stigeoclonium* (14.1%), followed by *Diatoma* (6.5%), *Gomphonema* (5.3%), *Nitzschia* (4.5%), and *Eunotia* (4.2%). Several of these genera had higher mean abundances during wet months, indicating that seasonal changes in flow, sediment delivery and nutrient inputs were associated with shifts in the relative contribution of dominant algal taxa.

**Table 8.** Main benthic algal genera contributing to Bray–Curtis dissimilarity between wet-and dry-season months based on SIMPER analysis. Mean wet-and dry-season densities are expressed as log10(cells m ² + 1). Contribution (%) indicates the percentage contribution of each genus to Bray–Curtis dissimilarity between seasons, and cumulative (%) indicates the running contribution of the listed genera to total assemblage dissimilarity. p-values are based on permutation tests.

| Genus | Mean density (Wet) | Mean density (Dry) | Contribution (%) | Cumulative (%) | *p* |
| --- | --- | --- | --- | --- | --- |
| *Lyngbya* | 6.80 | 6.17 | 19.41 | 19.41 | 0.0867 |
| *Stigeoclonium* | 6.60 | 6.15 | 14.12 | 33.53 | 0.9948 |
| *Diatoma* | 5.52 | 4.80 | 6.47 | 40.00 | 0.0015 |
| *Gomphonema* | 6.03 | 5.89 | 5.30 | 45.30 | 1.0000 |
| *Nitzschia* | 5.95 | 5.76 | 4.47 | 49.77 | 1.0000 |
| *Eunotia* | 5.33 | 4.70 | 4.15 | 53.92 | 0.0060 |
| *Navicula* | 5.70 | 5.63 | 3.94 | 57.86 | 0.9997 |
| *Oscillatoria* | 6.14 | 3.43 | 3.57 | 61.43 | 0.0011 |
| *Cyclotella* | 5.22 | 3.37 | 3.52 | 64.95 | 0.0001 |
| *Fragilaria* | 5.15 | 4.66 | 2.58 | 67.53 | 0.0279 |

### Environmental associations of benthic algal assemblage structure

Redundancy analysis (RDA) showed weak associations between benthic algal assemblage composition and measured physical and chemical variables across land-use categories (Fig. 7). The first two RDA axes accounted for 20.4% of the variation in assemblage composition, with RDA1 accounting for 12.3% and RDA2 for 8.1%. Environmental variables were screened for collinearity before interpretation. The final redundancy analysis retained seven environmental variables after removing variables with variance inflation factors (VIF) greater than 3. The retained variables were: turbidity, temperature, DO saturation, pH, NO□□–N, TP, SRP. The constrained component accounted for 31.68% of the total variation in algal community composition (R² = 0.317; Adjusted R² = 0.018). The overall RDA model was not statistically significant (p = 0.3693). Among the environmental variables, NO□□–N showed a significant marginal association with assemblage composition (*p* = 0.0239).

**Fig. 7.**
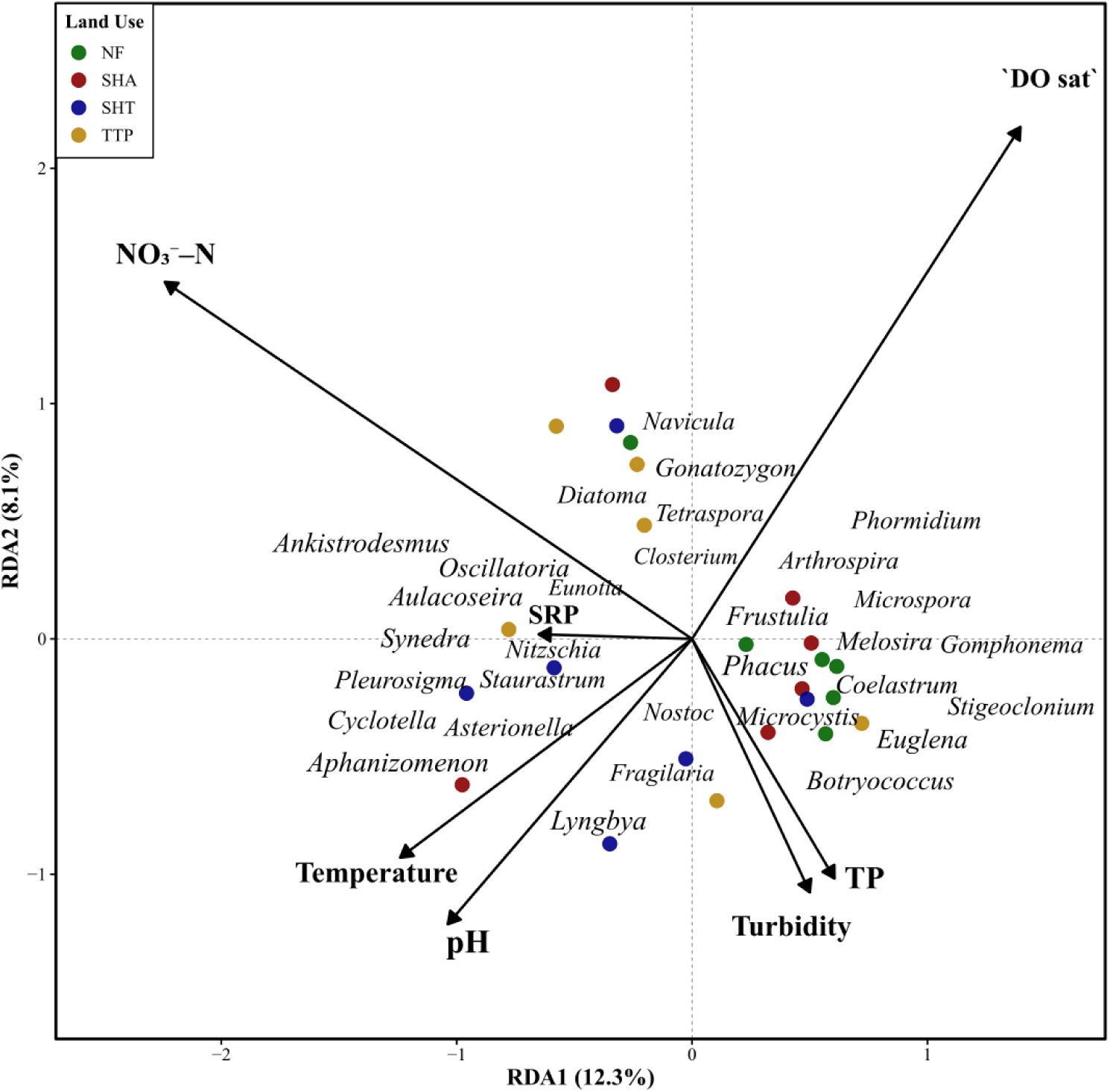
Redundancy analysis (RDA) ordination of algal community composition constrained by environmental variables. Sites are colored by land use type (NF = Natural Forest, SHA = smallholder agriculture, SHT = Smallholder Tea, TTP = Tea-tree Plantation). Environmental variables are shown as arrows. Genus labels are shown in italics

During monthly monitoring, the final RDA model retained temperature, turbidity, TP, DO, NO□□–N and TDS, following variance inflation factor (VIF) screening, with all retained variables exhibiting acceptable collinearity (Fig. 8). The constrained model explained 47.47% of the variation in benthic algal community composition (Adjusted R² = 0.325) and was statistically significant (global permutation test: F = 3.16, *p* = 0.031). The first canonical axis (RDA1), which accounted for 62.5% of the constrained variation, was significant (*p* = 0.030), whereas the second axis (RDA2) accounted for 20.4% of the constrained variation but was not significant (*p* = 0.464). Permutation tests indicated that temperature was the only environmental variable significantly associated with variation in algal community composition, whereas turbidity, TP, DO, NO□□–N and TDS did not explain significant additional variation after accounting for the overall model. Algal taxa aligned in the direction of the environmental vectors showed associations with the measured gradients; for example, *Lyngbya, Gomphonema,* and *Euglena* were associated with the temperature and TDS gradients, whereas *Eunotia* and *Nostoc* were more closely associated with higher dissolved oxygen.

**Fig. 8.**
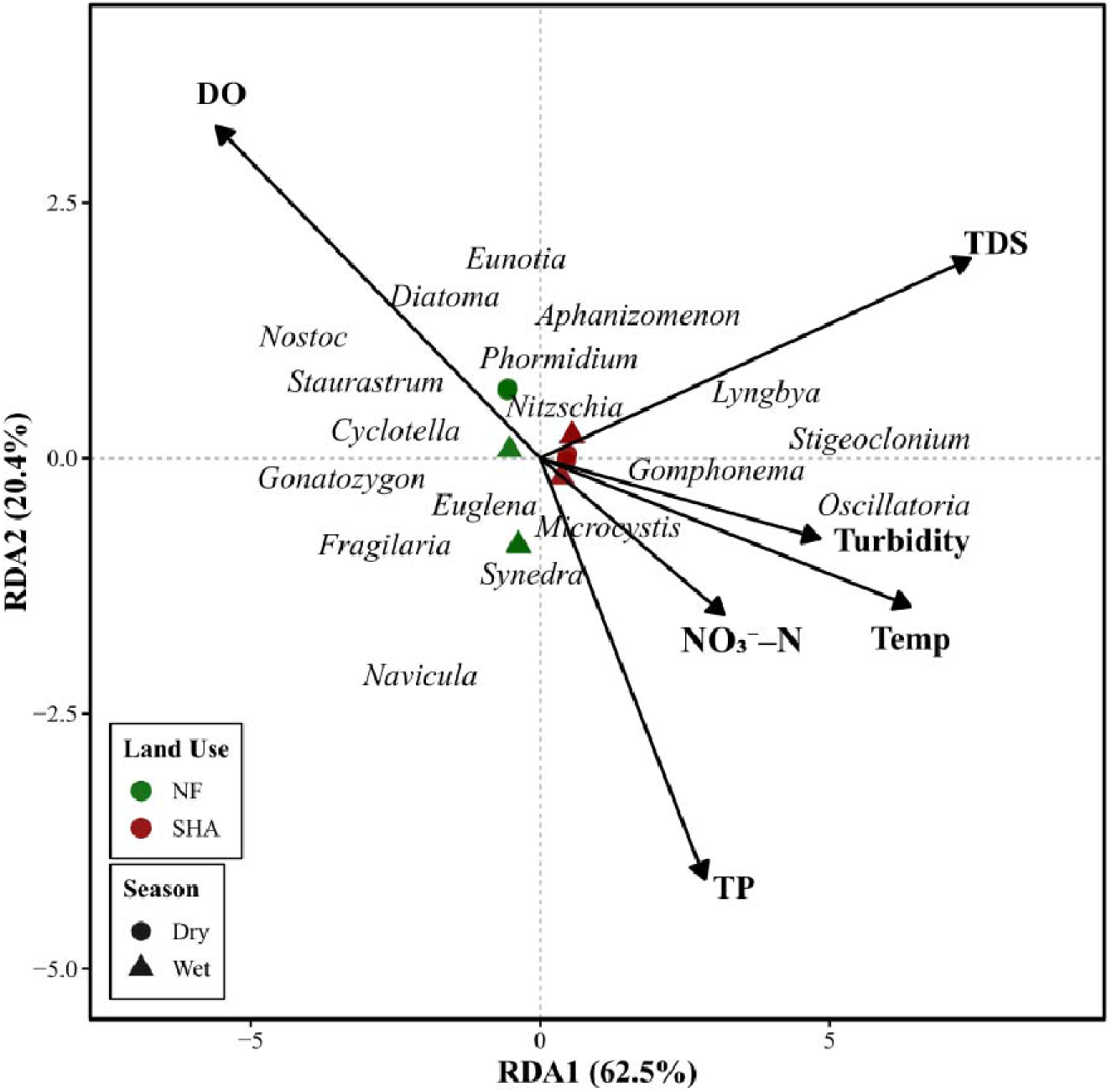
Redundancy analysis (RDA) showing associations between benthic algal assemblage composition and environmental variables for monthly monitoring samples from representative natural forest (NF) and smallholder agriculture (SHA) streams. Colours indicate land use, symbols indicate season. Environmental variables are shown as arrows. Genus labels are shown in italics

## Discussion

This study examined how catchment land-use and seasonality influence benthic algal assemblages and biofilm biomass in headwater streams of the Sondu–Miriu River Basin. By integrating a synoptic survey with temporal monitoring at two streams (NF-1 and SHA-14), the study captured spatial variation among land-use categories and temporal variation within the two monitored streams. Land use was associated with significant differences in electrical conductivity, total dissolved solids, dissolved oxygen and temperature, while turbidity showed a higher but non-significant mean in the agriculturally influenced streams. Studies from tropical and subtropical catchments have reported similar differences, where streams draining forested landscapes typically exhibit better water quality (e.g., higher dissolved oxygen and lower turbidity) than those draining agricultural or disturbed catchments (de Mello et al., 2018; Hirko et al., 2025; Uriarte et al., 2011). Wet–dry variation also influenced these spatial patterns by altering sediment transport, organic matter inputs, and nutrient delivery. In many tropical headwater systems, rainfall-driven runoff and changes in discharge generate strong seasonal fluctuations in water quality and biological communities, often amplifying land-use signals during high-flow periods (Rodrigues et al., 2018; Jerves-Cobo et al., 2020; Dalu et al., 2022). These interacting spatial and temporal gradients were reflected in benthic algal assemblages in the present study. Forest streams supported more diverse and evenly distributed communities, whereas agriculturally influenced streams showed stronger dominance patterns and greater accumulation of organic biofilm material. These findings indicate that land-use provides the primary environmental template structuring algal assemblages, while seasonal variability regulates the magnitude and short-term dynamics of these ecological responses.

### Water physical and chemical variables

Physical variables that differed significantly across land-use categories (EC, TDS, DO and temperature) collectively pointed to systematic contrasts consistent with differing degrees and types of catchment modification. NF streams exhibited the highest dissolved oxygen, which agrees with Minaya et al. (2013) and Masese et al. (2017), who reported lower DO concentrations in agricultural streams compared to their forested counterparts, aligning with comparatively better oxygenation in less disturbed reaches. In contrast, SHA streams showed the highest EC and TDS, indicating greater dissolved ionic content in agriculturally influenced streams (Jacobs et al., 2017; Masese et al., 2017; Wanderi et al., 2022). Notably, SHT and SHA streams recorded higher turbidity, suggesting that these streams experience greater input of fine sediments from eroding cultivated slopes, footpaths, drainage channels, and streambank disturbance. The higher temperature in SHT streams may reflect reduced riparian shading and greater exposure of the stream channel to direct solar radiation (Stenfert Kroese et al., 2020) relative to the other land-use settings (Table 1). Similar physical and chemical gradients and land-use-associated contrasts have been reported in other tropical and Afromontane catchments, where disturbance alters sediment delivery, solute loads, and oxygen regimes (e.g., Jacobs et al., 2017; Stenfert Kroese et al., 2020; Masese et al., 2024).

Seasonal monitoring further indicated that some of these physical and chemical variables fluctuated over time, especially during periods of increased rainfall and discharge (Table 2). Variables such as turbidity, suspended sediments, and organic matter showed marked increases during high-flow conditions, reflecting rainfall-driven mobilization of fine sediments and organic material from surrounding catchments (Kuluo et al., 2026; Stenfert Kroese et al., 2020; Jacobs et al., 2018). These temporal fluctuations suggest that while land use establishes persistent physical and chemical gradients across streams, seasonal hydrological processes can amplify or modulate these gradients through changes in sediment transport, runoff connectivity, and organic matter delivery (Rodrigues et al., 2018). The seasonal response of nitrite observed in the monthly monitoring data is consistent with this interpretation, as NO□□–N is a transient form of inorganic nitrogen whose concentration can change with runoff inputs, oxygen availability and microbial nitrogen transformations in stream water and sediments (Peterson et al., 2001; Bernot & Dodds, 2005; Garnier et al., 2009).

Elevated turbidity and TSS observed in SHT and SHA streams are consistent with enhanced mobilization and transport of fine particulates, which can arise from cultivated lands, pathway connectivity (e.g., human and livestock tracks, drains), and bank or instream disturbance and erosion caused by human and livestock activity (Defersha & Melesse, 2012; Stenfert Kroese et al., 2020; Iteba et al., 2021). However, high variability observed in SHT turbidity and TSS during the synoptic survey reflects the episodic nature of sediment transport in these headwater catchments. While the synoptic sampling in August captured elevated TSS, single-point sampling often fails to capture the maximum sediment peaks associated with high-intensity rainfall events. This limitation is evident when comparing the synoptic results to our monthly monitoring data; during the wet season, the monitored SHA-14 site recorded higher mean TSS levels, which were higher than the broader synoptic average. These results are consistent with previous studies in the region (Jacobs et al., 2018; Stenfert Kroese et al., 2020) which show that agricultural and tea-growing landscapes are major sources of episodic sediment delivery driven by short-lived runoff pulses from cultivated slopes and livestock tracks. Higher EC and TDS in SHA streams similarly align with increased delivery of dissolved ions from agricultural areas and human settlements, while the comparatively higher DO in NF is consistent with reduced organic loading and reduced human disturbance. The lower DO observed in SHT streams is consistent with higher temperature and greater inputs of fine sediment and organic matter, which can increase microbial respiration and sediment oxygen demand. Low DO often reflects high system oxygen demand (microbial decomposition and community respiration) and reduced reaeration, conditions that can occur where fine sediments accumulate, biofilms thicken, and organic matter inputs increase (Piatka et al., 2022; dos Reis Oliveira et al., 2019; Hirko et al., 2025).

Nutrient peaks observed during the wet season coincided with elevated turbidity, suspended sediments and particulate organic matter, suggesting that rainfall-driven runoff enhanced the mobilization and transport of organic matter and nutrients from surrounding catchments into stream channels. Similar seasonal pulses in nutrient delivery have been documented in tropical headwater systems, where increased wet-season hydrological connectivity enhances the transfer of nutrients and organic matter from terrestrial to aquatic environments (Jacobs et al., 2018; Masese et al., 2024).

### Benthic algal assemblage structure

Relative-abundance patterns indicate that land-use modification altered streambed conditions in ways that favoured dominance by a limited set of taxa and reduced assemblage evenness (Hart et al., 2013). The pronounced dominance of *Stigeoclonium* in SHT streams may reflect the ability of disturbance-tolerant filamentous algae to persist under warmer, more turbid and potentially less stable stream conditions (Stancheva & Sheath, 2016). Comparable shifts have been reported in tropical mountain streams where riparian degradation and open land use altered periphyton composition, abundance, and diversity, and in-stream benthic biofilms exposed to anthropogenic land-use, where community structure and function changed in response to altered environmental conditions (Cartuche et al., 2026; Qu et al., 2017). Seasonal monitoring indicates that these spatial patterns were superimposed on temporal turnover: high-flow periods likely disturbed established biofilms and promoted recolonization, whereas lower-flow periods allowed recovery and stronger expression of dominant taxa. This interpretation is consistent with evidence from tropical streams showing that storm flows can strongly reduce epilithic biomass and favour resistant or rapidly colonizing algal forms (Schneider & Petrin, 2017).

The marked differences in diversity, dominance, and biomass among land-use types indicate that disturbance acted primarily through community reorganization (Casartelli & Ferragut, 2018). NF streams supported the most even assemblages, and were characterised mainly by diatom genera, including *Diatoma*, *Cyclotella*, and *Eunotia*. These genera were more common in the less disturbed forested streams, where turbidity, EC, TDS, and temperature were lower and dissolved oxygen was higher. In contrast, SHA streams were dominated mainly by *Lyngbya* spp., with additional contributions from *Stigeoclonium* and *Microcystis*, while SHT streams were dominated by *Stigeoclonium*, with smaller contributions from *Phormidium*, *Synedra*, and *Melosira*. These taxa were more common in streams with higher turbidity, temperature, EC, and TDS. This suggests that land-use effects were expressed more strongly through changes in the relative abundance of dominant genera (Zhao et al., 2023). Similar patterns have been reported from tropical headwater streams, where local environmental conditions explained biofilm diversity more strongly than broad land-use categories alone, and from tropical mountain streams where degraded reaches supported the most divergent periphyton assemblages (Burgos-Caraballo et al., 2014; Cartuche et al., 2026). The monthly monitoring data showed a similar pattern. The NF stream remained diatom-rich in both seasons, with *Diatoma*, *Eunotia*, *Cyclotella*, *Fragilaria*, *Navicula* and *Nitzschia* contributing most of the assemblage. In contrast, the SHA stream was characterized by higher relative abundances of *Lyngbya* spp. and the filamentous green alga *Stigeoclonium* in both seasons, with additional contributions from *Oscillatoria*, *Gomphonema*, *Nitzschia* and *Microcystis* (Burgos-Caraballo et al., 2014).

Biomass proxies provided insight into how land-use influenced benthic biofilm structure and function (Qu et al., 2017). Synoptic patterns showed that chlorophyll-*a* and AFDM were highest in SHA streams. Chlorophyll-a indicated pigment-based autotrophic biomass, whereas AFDM represented total organic biofilm material on benthic substrates (Biggs & Kilroy, 2000; Myers, 2014). AFDM gave the clearer signal of agricultural influence, with the highest values recorded in SHA streams, indicating greater organic biofilm accumulation in these reaches. Higher AFDM in the SHA stream suggests greater organic biofilm accumulation under lower-flow conditions. Autotrophic Index described the amount of total organic biofilm material relative to chlorophyll-a and therefore complemented the separate AFDM and chlorophyll-a measurements (Myers, 2014; Baattrup-Pedersen et al., 2020).

Monthly monitoring showed similar patterns. Chlorophyll-a and AFDM were higher during dry months in both monitoring streams, with the highest AFDM recorded in the SHA stream. Higher AFDM in the SHA stream suggests greater accumulation of benthic organic material in agriculturally influenced reaches, particularly under lower-flow conditions. Similar dry-season increases in benthic organic matter and periphyton biomass have been reported in stream studies where reduced discharge favoured biofilm accumulation and stability on stream substrates (Biggs & Kilroy, 2000; Beck et al., 2019; Taniwaki et al., 2019). The autotrophic index supported this interpretation as an AFDM/chlorophyll-a ratio, describing total organic biofilm material relative to algal pigment. Comparable seasonal changes in periphyton biomass have been linked to changes in flow conditions, sediment deposition, nutrient availability, and organic matter inputs in lotic systems (Biggs & Kilroy, 2000; Huang et al., 2018).

### Ordination patterns and environmental associations of benthic algal assemblages

Multivariate ordination analyses (NMDS, PERMANOVA) showed significant differences in benthic algal assemblage composition among land-use categories, whereas the synoptic RDA indicated only weak, non-significant associations between assemblage composition and the measured physical and chemical variables. In the synoptic dataset, the ordination suggested that NF assemblages aligned with higher DO and were associated with diatom-dominated taxa such as *Diatoma*, *Cyclotella*, and *Eunotia* exhibiting higher diversity and evenness. Elevated dissolved oxygen is commonly linked to lower organic loading, greater habitat stability, and reduced chronic disturbance, conditions that favour coexistence among multiple benthic algal taxa without strong dominance by a single genus. This pattern supports the interpretation that physical and chemical conditions characteristic of forested catchments promote structurally complex algal assemblages. In contrast, SHA streams occurred closer to higher TDS, turbidity and water temperature, and aligned with assemblages characterized by elevated AFDM and more organic-rich biofilms, indicating a dissolved-constituent and organic-matter regime typical of agriculturally influenced streams (Masese et al., 2017; de Mello et al., 2018). SHT streams also occurred closer to higher temperature and turbidity and clustered with filamentous genera, particularly *Stigeoclonium*. This association may reflect the occurrence of filamentous forms under more turbid or physically disturbed streambed conditions. The low chlorophyll-a concentrations observed in SHT streams suggest that structural dominance in relative abundance did not necessarily correspond to higher pigment-based autotrophic biomass, because dominance reflects proportional composition whereas chlorophyll-a represents absolute pigment mass per unit area (Morris et al., 2014; Myers, 2014).

For the monthly monitoring dataset, samples from the SHA stream aligned more closely with higher temperature, TDS, TP, nitrate, and were associated with genera such as *Lyngbya*, *Stigeoclonium*, *Gomphonema* and *Euglena*. This pattern is consistent with the assemblage results, where the SHA stream supported higher relative abundances of filamentous green algae and cyanobacteria during monitoring. In the monthly RDA, temperature was the only retained variable significantly associated with algal assemblage composition. However, the small temperature differences among monitored samples suggest that this association may reflect broader environmental differences between the NF and SHA streams, with limited evidence for an independent temperature effect. Stream algae have short generation times and can respond rapidly to changes in light, flow, nutrient availability and streambed conditions (Stevenson & Rollins, 2017; Tromboni et al., 2019; Penk et al., 2024). In contrast, NF samples were positioned closer to higher dissolved oxygen conditions and were associated with diatom-rich assemblages. This pattern is consistent with the higher oxygen conditions observed in the NF stream and with the persistence of diatom-rich assemblages across the monitoring period.

### Benthic algae as indicators of land use disturbance

A key implication of this study is that benthic algae responded to land-use disturbance through changes in assemblage composition, diversity, and relative abundance of dominant genera. Shannon and Simpson diversity differed among land-use categories, while total algal density showed weaker separation. These differences were mainly explained by changes in the relative abundance of dominant genera. NF streams supported more diatom-rich assemblages, while agriculturally influenced streams showed greater representation of filamentous green algae and cyanobacteria. These compositional shifts show how catchment modification was reflected in benthic algal community structure, complementing the physical and chemical patterns. Similar responses of benthic algal and diatom assemblages to agricultural and catchment disturbance have been reported in other stream systems (Pan et al., 1996; Stevenson & Rollins, 2017; Bere & Tundisi, 2011).

Biomass proxies complemented the assemblage results because benthic chlorophyll-a, AFDM, and autotrophic index captured different aspects of benthic algal and organic matter accumulation across land-use categories and sampling months. AFDM showed the strongest and most consistent response to land-use particularly in SHA, indicating greater accumulation of organic biofilm material in agricultural streams. Because chlorophyll-a reflects autotrophic biomass whereas AFDM integrates total organic biofilm mass, their divergence provides complementary information on biofilm condition and functioning (Myers, 2014). In contrast, the Autotrophic Index differed among land-use categories and also showed marked wet–dry variation within the two monitored streams (Biggs & Kilroy, 2000; Stevenson & Rollins, 2017; Huang et al., 2018).

The SIMPER results support the relative abundance and ordination findings by showing that land-use differences were associated with changes in dominant genera, especially filamentous green algae, cyanobacteria and selected diatoms. Together with the multivariate analyses, which showed compositional differences among land-use categories but only weak environmental associations in the synoptic RDA, these findings support a multi-metric framework in which diversity/dominance metrics such as diversity indices, and biomass metrics such as AFDM are considered for detecting land-use impacts, while chlorophyll-a provides a complementary measure of autotrophic biomass and the Autotrophic Index describes wet–dry differences in total organic biofilm material relative to algal pigment within the two monitored streams (Weber, 1973; Myers, 2014).

## Conclusion

This study showed that benthic algal assemblage composition and periphyton biomass metrics varied among the four land-use categories across the 24 sampled streams, while monthly monitoring of one NF stream and one SHA stream showed additional descriptive wet–dry variation. Streams draining natural forest supported more even, diatom-rich algal assemblages, whereas smallholder agriculture and smallholder tea streams showed stronger dominance by a few genera, including filamentous green algae and cyanobacteria. These assemblage shifts were accompanied by differences in water quality, with natural forest streams showing higher dissolved oxygen and agriculturally influenced streams showing higher electrical conductivity, total dissolved solids, turbidity or temperature. Tea–tree plantation streams showed intermediate values for several physical and chemical variables and biomass measures, while their assemblage composition did not differ significantly from natural forest streams in the pairwise PERMANOVA comparison. In the two monitored streams, wet-season months were associated with higher suspended sediments, particulate organic matter and nutrient concentrations, especially in the smallholder agriculture stream.

Periphyton biomass proxies helped distinguish different aspects of biofilm condition. Ash-free dry mass was particularly useful for showing organic biofilm accumulation in agriculturally influenced streams. Chlorophyll-a described pigment-based autotrophic biomass, while the autotrophic index described the amount of total organic biofilm material relative to algal pigment. The findings support the use of benthic algal assemblage composition together with periphyton biomass proxies, especially ash-free dry mass, for assessing land-use effects in Afrotropical headwater streams. Future monitoring should combine algal assemblage data, biomass proxies and repeated seasonal sampling, particularly during wet periods when sediment and nutrient inputs are likely to increase. This would strengthen the use of benthic algae in routine ecological assessment and improve detection of land-use impacts in tropical headwater catchments.

## Statements and Declarations

## Acknowledgements

We acknowledge the Department of Fisheries and Aquatic Sciences, University of Eldoret, for providing laboratory space and equipment used during this work. We are grateful to Christian Bashizi, Kelvin Moenga, Ruto Vincent, and Lubanga Henry for their valuable assistance in field sampling and laboratory analysis. Finally, we extend our appreciation to colleagues and friends whose insights and encouragement contributed to the successful completion of this study.

## Author contributions

SNR: Conceptualization, formal analysis, writing-original draft, and visualization; FM: Conceptualization, methodology, validation, supervision; FM, SJ, and LB: Project administration and funding acquisition; SNR, JI, GK, SJ and FM: investigation; SNR, JI and GK: Data curation. All authors commented on the manuscript and approved it for submission.

## Data availability

Data and R code are available from the corresponding author upon request.

